# Predicting microbiome compositions from species assemblages through deep learning

**DOI:** 10.1101/2021.06.17.448886

**Authors:** Sebastian Michel-Mata, Xu-Wen Wang, Yang-Yu Liu, Marco Tulio Angulo

**Affiliations:** Center for Applied Physics and Advanced Technology, Universidad Nacional Autónoma de México, Juriquilla 76230, México; Department of Ecology and Evolutionary Biology, Princeton University, Princeton, NJ 08544, USA; Channing Division of Network Medicine, Department of Medicine, Brigham and Women’s Hospital and Harvard Medical School, Boston, Massachusetts 02115, USA; CONACyT - Institute of Mathematics, Universidad Nacional Autónoma de México, Juriquilla 76230, México

## Abstract

Microbes can form complex communities that perform critical functions in maintaining the integrity of their environment or their hosts’ well-being. Rationally managing these microbial communities requires improving our ability to predict how different species assemblages affect the final species composition of the community. However, making such a prediction remains challenging because of our limited knowledge of the diverse physical, biochemical, and ecological processes governing microbial dynamics. To overcome this challenge, here we present a deep learning framework that automatically learns the map between species assemblages and community compositions from training data only, without knowledge of any of the above processes. First, we systematically validate our framework using synthetic data generated by classical population dynamics models. Then, we apply it to experimental data of both *in vitro* and *in vivo* communities, including ocean and soil microbial communities, *Drosophila melanogaster* gut microbiota, and human gut and oral microbiota. In particular, we show how our framework learns to perform accurate out-of-sample predictions of complex community compositions from a small number of training samples. Our results demonstrate how deep learning can enable us to understand better and potentially manage complex microbial communities.

## Introduction

Microbes can form complex multispecies communities that perform critical functions in maintaining the integrity of their environment^1,2^ or the well-being of their hosts^3–6^. For example, microbial communities play key roles in nutrient cycling in soils^7^ and crop growth^8^. In humans, the gut microbiota plays important roles in our nutrition^9^, immune system response^10^, pathogen resistance^11^, and even our nervous central system response^5^. Still, species invasions (e.g., pathogens) and extinctions (e.g., due to antibiotic administration) produce changes in the species assemblages that may shift these communities to undesired compositions^12^. For instance, antibiotic administrations can shift the human gut microbiota to compositions making the host more susceptible to recurrent infections by pathogens^13^. Similarly, intentional changes in the species assemblages, such as by using fecal microbiota transplantations, can shift back these communities to desired “healthier” compositions^14,15^. Therefore, improving our ability to rationally manage these microbial communities requires that we can predict changes in the community composition based on changes in species assemblages^16^. Building these predictions would also reduce managing costs, helping us to predict which changes in the species’ assemblages are more likely to yield a desired community composition. Unfortunately, making such a prediction remains challenging because of our limited knowledge of the diverse physical^17^, biochemical^18^, and ecological^19,20^ processes governing the microbial dynamics.

To overcome the above challenge, here we present a deep learning framework that automatically learns the map between species assemblages and community compositions from training data only, without knowledge of the underlying microbial dynamics. We systematically validated our framework using synthetic data generated by classical ecological dynamics models, demonstrating its robustness to changes in the system dynamics and to measurement errors. Then, we applied our framework to experimental data of both *in vitro* and *in vivo* communities, including ocean and soil microbial communities^21,22^, *Drosophila melanogaster* gut microbiota^23^, and human gut and oral microbiota^24^. Across these diverse microbial communities, we show how our framework learns to predict accurate out-of-sample compositions given a few training samples. Our results show how deep learning can be an enabling ingredient for understanding and managing complex microbial communities.

## Methods

Consider the pool *Ω* = {1, …, *N* } of all microbial species that can inhabit an ecological habitat of interest, such as the human gut. A microbiome sample obtained from this habitat can be considered as a local community assembled from *Ω* with a particular *species assemblage*. The species assemblage of a sample is characterized by a binary vector *z* ∈ {0, 1}^*N*^, where its *i*-th entry *z*_*i*_ satisfies *z*_*i*_ = 1 (or *z*_*i*_ = 0) if the *i*-th species is present (or absent) in this sample. Each sample is also associated with a *composition* vector *p* ∈ *Δ*^*N*^, where *p*_*i*_ is the relative abundance of the *i*-th species, and 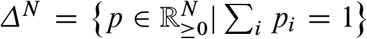 is the probability simplex. Mathematically, our problem is to learn the map

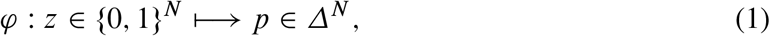

which assigns the composition vector *p* = *φ*(*z*) based on the species assemblage *z*.

Knowing the above map would be instrumental in understanding the assembly rules of microbial communities^25^. However, learning this map is a fundamental challenge because the map depends on many physical, biochemical, and ecological processes influencing the dynamics of microbial communities. These processes include the spatial structure of the ecological habitat^17^, the chemical gradients of available resources^18^, and inter/intra-species interactions^20^, to name a few. For large microbial communities like the human gut microbiota, our knowledge of all these processes is still rudimentary, hindering our ability to predict microbial compositions from species assemblages.

Next, we show it is possible to predict the microbial composition from species assemblage without knowing the mechanistic details of the above processes. Our solution is a deep learning framework that learns the map *φ* directly from a dataset 𝔇 of *S* samples, each of which is associated with a pair (*z, p*), see Fig. 1.

**Figure 1:**
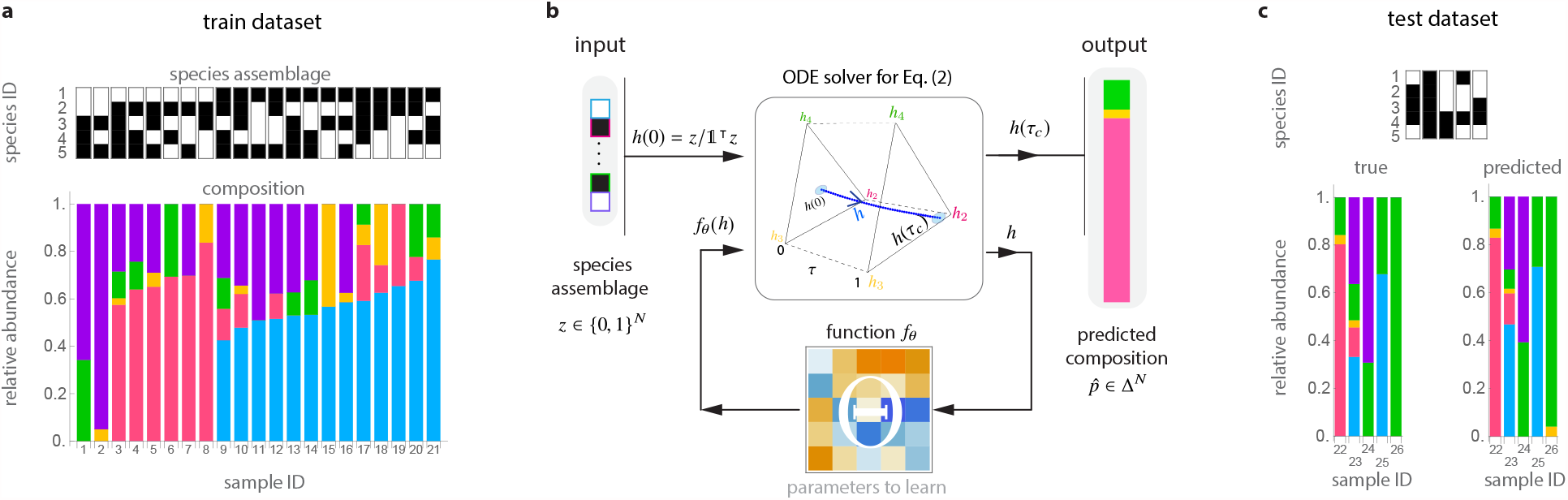
A deep learning framework to predict microbiome compositions from species assemblages. We illustrate this framework using experimental data from a pool of *N* = 5 bacterial species in *Drosophila melangaster* gut microbiota^23^: *Lactobacillus plantarum* (blue), *Lactobacillus brevis* (pink), *Acetobacter pasteurianus* (yellow), *Acetobacter tropicalis* (green), and *Acetobacter orientalis* (purple). **a**. We randomly split this dataset into training (𝔇_1_) and test (𝔇_2_) datasets, which contain 80% and 20% of the samples, respectively. Each dataset contains pairs (*z, p*) with the species assamblage *z* ∈ {0, 1}^*N*^ (top) and its corresponding composition *p* ∈ *Δ*^*N*^ (bottom) from each sample. **b**. To predict compositions from species assamblages, our cNODE framework consists of a solver for the ODE shown in Eq. (2), together with a chosen parametrized function *f*_*θ*_. During training, the parameters *θ* are adjusted to learn to predict the composition 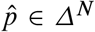 of the species assamblage *z* ∈ {0, 1}^*N*^ in 𝔇_1_. **c**. After training, the performance is evaluated by predicting the composition of never-seen-before species assamblages in the test dataset 𝔇_2_. In this experimental microbiota, cNODE learned to perform accurate predictions of the composition in the test dataset. For example, in the assemblage of species 3 and 4 (sample 26), cNODE correctly predicts that the composition is strongly dominated by a single species.

### Conditions for predicting compositions from species assemblages

To ensure that the problem of learning *φ* from 𝔇 is mathematically well-posed, we make the following assumptions. *First*, we assume that the species pool in the habitat has universal dynamics^26^ (i.e., different local communities of this habitat can be described by the same population dynamics model with the same parameters). This assumption is necessary because, otherwise, the map *φ* does not exist, implying that predicting community compositions from species assemblages has to be done in a sample-specific manner, which is a daunting task. For *in vitro* communities, this assumption is satisfied if samples were collected from the same experiment or multiple experiments but with very similar environmental conditions. For *in vivo* communities, empirical evidence indicates that the human gut and oral microbiota of healthy adults, as well as certain environment microbiota, display strong universal dynamics^26^. *Second*, we assume that the compositions of those collected samples represent steady states of the microbial communities. This assumption is natural because the map *φ* is not well defined for highly fluctuating microbial compositions. We note that observational studies of host-associated microbial communities such as the human gut microbiota indicate that they remain close to stable steady states in the absence of drastic dietary change or antibiotic administrations^24,27,28^. *Finally*, we assume that for each species assemblage *z* ∈ {0, 1}^*N*^ there is a unique steady-state composition *p* ∈ *Δ*^*N*^. In particular, this assumption requires that the true multi-stability does not exist for the species pool (or any subset of it) in this habitat. This assumption is required because, otherwise, the map *φ* is not injective, and the prediction of community compositions becomes mathematically ill-defined. In practice, we expect that the above three assumptions cannot be strictly satisfied. Therefore, any algorithm that predicts microbial compositions from species assemblages needs to be systematically tested to ensure its robustness against errors due to the violation of such approximations.

### Limitations of traditional deep learning frameworks

Under the above assumptions, a straightforward approach to learning the map *φ* from 𝔇 would be using deep neural networks^29,30^ such as a feedforward Residual Network^31^ (ResNet). As a top-rated tool in image processing, ResNet is a cascade of *L* ≥ 1 hidden layers where the state *h*_𝓁_ ∈ ℝ^*N*^ of the 𝓁-th hidden layer satisfies *h*_𝓁_ = *h*_𝓁 − 1_ + *f*_*θ*_ (*h*_𝓁 − 1_), 𝓁 = 1, … *L*, for some parametrized function *f*_*θ*_ with parameters *θ*. These hidden layers are plugged to the input *h*_0_ = *g*_in_(*z*) and the output 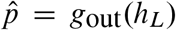 layers, where *g*_in_ and *g*_out_ are some functions. Crucially, for our problem, any architecture must satisfy two restrictions: (1) vector 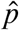 must be compositional (i.e., 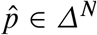); and (2) the predicted relative abundance of any absent species must be identically zero (i.e., *z*_*i*_ = 0 should imply that 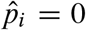). Simultaneously satisfying both restrictions requires that the output layer is a normalization of the form 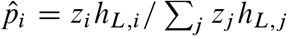, and that *f*_*θ*_ is a non-negative function (because *h*_*L*_ ≥ 0 is required to ensure the normalization is correct). We found that it is possible to train such a ResNet for predicting compositions in simple cases like small *in vitro* communities (Supplementary Note S2.1). But for large *in vivo* communities like the human gut microbiota, ResNet does not perform very well (Supplementary Fig. S1). This result is likely due to the normalization of the output layer, which challenges the training of neural networks because of vanishing gradients^30^.

The vanishing gradient problem is often solved by using other normalization layers such as the `softmax` or `sparsemax` layers^32^. However, we cannot use these layers because they do not satisfy the second restriction. We also note that ResNet becomes a universal approximation only in the limit *L* → ∞, which again complicates the training^33^.

### A new deep learning framework

To overcome the limitations of traditional deep learning frameworks based on neural networks (such as ResNet) in predicting microbial compositions from species assemblages, we developed cNODE (compositional Neural Ordinary Differential Equation), see Fig. 1b. The cNODE framework is based on the notion of Neural Ordinary Differential Equations, which can be interpreted as a continuous limit of ResNet where the hidden layers *h*’s are replaced by an ordinary differential equation (ODE)^34^. In cNODE, an input species assemblage *z* ∈ {0, 1}^*N*^ is first transformed into the initial condition *h*(0) = *z* / 𝕝^⊤^ *z* ∈ *Δ*^*N*^, where 𝕝 = (1, …, 1)^⊤^ ∈ ℝ^*N*^ (left in Fig. 1b). This initial condition is used to solve the set of nonlinear ODEs

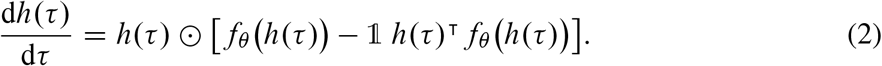

Here, the independent variable *τ* ≥ 0 represents a virtual “time”. The expression *h* ⊙ *v* is the entry-wise multiplication of the vectors *h, v* ∈ ℝ^*N*^. The function *f*_*θ*_ : *Δ*^*N*^ → ℝ^*N*^ can be any continuous function parametrized by *θ*. For example, it can be the linear function *f*_*θ*_ (*h*) = *Θh* with parameter matrix *Θ* ∈ ℝ^*N*×*N*^ (bottom in Fig. 1b), or a more complicated function represented by a feedforward deep neural network. Note that Eq. (2) is a general form of the replicator equation —a canonical model in evolutionary game theory^35^— with *f*_*θ*_ representing the fitness function. By choosing a final integration “time” *τ*_*c*_ > 0, Eq. (2) is numerically integrated to obtain the prediction 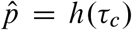 that is the output of cNODE (right in Fig. 1b). We choose *τ*_*c*_ = 1 without loss of generality, as *τ* in Eq. (2) can be rescaled by multiplying *f*_*θ*_ by a constant. The cNODE thus implements the map

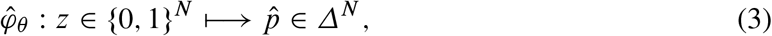

taking an input species assemblage *z* to the predicted composition 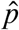 (see Supplementary Note S1 for implementation details). Note that Eq. (2) is key to cNODE because its architecture guarantees that the two restrictions imposed before are naturally satisfied. Namely, 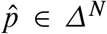 because the conditions *h*(0) ∈ *Δ*^*N*^ and 𝕝^⊤^d*h* / d*τ* = 0 imply that *h*(*τ*) ∈ *Δ*^*N*^ for all *τ* ≥ 0. Additionally, *z*_*i*_ = 0 implies 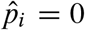 because *h*(0) and *z* have the same zero pattern, and the right-hand side of Eq. (2) is entry-wise multiplied by *h*.

We train cNODE by adjusting the parameters *θ* to approximate *φ* with 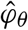. To do this, we first choose a distance or dissimilarity function *d*(*p, q*) to quantify how dissimilar are two compositions *p, q* ∈ *Δ*^*N*^. One can use any Minkowski distance or dissimilarity function. In the rest of this paper, we choose the Bray-Curtis^36^ dissimilarity to present our results. Specifically, for a dataset 𝔇_*i*_ ⊆ 𝔇, we use the loss function

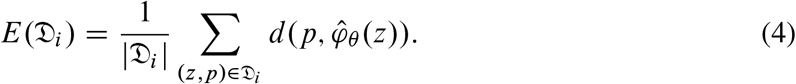

Second, we randomly split the dataset 𝔇 into training 𝔇_1_ and test 𝔇_2_ datasets. Next, we choose an adequate functional form for *f*_*θ*_. In our experiments, we found that the linear function *f*_*θ*_ (*h*) = *Θh, Θ* ∈ ℝ^*N*×*N*^, provides accurate predictions for the composition of *in silico, in vitro*, and *in vivo* communities. Despite *f*_*θ*_ is linear, the map 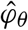 is nonlinear because it is the solution of the nonlinear ODE of Eq. (2). Finally, we adjust the parameters *θ* by minimizing Eq. (4) on 𝔇_1_ using a gradient-based meta-learning algorithm^37^. This learning algorithm enhances the generalizability of cNODE (Supplementary Note S1.2 and Supplementary Fig. S1). Once trained, we calculate cNODE’s test prediction error *E*(𝔇_2_) that quantifies cNODE’s performance in predicting the compositions of never-seen-before species assemblages. Test prediction errors could be due to a poor adjustment of the parameters (i.e., inaccurate prediction of the training set), low ability to generalize (i.e., inaccurate predictions of the test dataset), or violations of our three assumptions (universal dynamics, steady-state samples, no true multi-stability).

Figure 1 shows the result of applying cNODE to fly gut microbiome samples collected in an experimental study^23^. In this study, germ-free flies (*Drosophila melanogaster*) were colonized with all possible combinations of *N* = 5 core species of fly gut bacteria, i.e., *Lactobacillus plantarum* (species-1), *Lactobacillus brevis* (species-2), *Acetobacter pasteurianus* (species-3), *Acetobacter tropicalis* (species-4), and *Acetobacter orientalis* (species-5). The dataset contains 41 replicates for the composition of each of the 2^*N*^ − 1 = 31 local communities with different species assamblages. To apply cNODE, we aggregated all replicates and calculated their average composition, resulting in one “representative” sample per species assamblage (Supplementary Note S4). We also excluded the trivial samples with a single species, resulting in *S* = 26 samples. We trained cNODE by randomly choosing 21 of those samples (80%) as the training dataset (Fig. 1a). Once trained, cNODE accurately predicts microbial compositions in the test dataset of 5 species assemblages (Fig. 1c). For example, cNODE predicts that in the assemblage of species 3 with species 4, species 3 will become nearly extinct, which agrees well with the experimental result (sample 26 in Fig. 1c).

## Results

### *In silico* validation of cNODE

To systematically evaluate the performance cNODE, we generated *in silico* data for pools of *N* = 100 species with population dynamics given by the classic Generalized Lotka-Volterra (GLV) model^38^

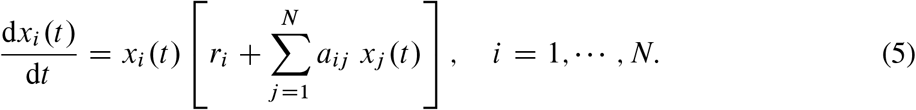

Above, *x*_*i*_ (*t*) denotes the abundance of the *i*-th species at time *t* ≥ 0. The GLV model has as parameters the interaction matrix *A* = (*a*_*ij*_) ∈ ℝ^*N* × *N*^, and the intrinsic growth-rate vector *r* = (*r*_*i*_) ∈ ℝ^*N*^. The parameter *a*_*ij*_ denotes the inter- (if *j* ≠ *i*) or intra- (if *j* = *i*) species *interaction strength* of species *j* to the per-capita growth rate of species *i*. The parameter *r*_*i*_ is the intrinsic growth rate of species *i*. The interaction matrix *A* determines the ecological network *𝒢*(*A*) underlying the species pool. Namely, this network has one node per species and edges (*j* → *i*) ∈ *𝒢*(*A*) if *a*_*ij*_ ≠ 0. The *connectivity C* ∈ [0, 1] of this network is the proportion of edges it has compared to the *N* ^2^ edges in a complete network. Despite its simplicity, the GLV model successfully describes the population dynamics of microbial communities in diverse environments, from the soil^39^ and lakes^40^ to the human gut^11,41,42^. To validate cNODE, we generated synthetic microbiome samples as steady-state compositions of GLV models with random parameters by choosing *a*_*ij*_ ∼ Bernoulli (*C*) Normal (0, *σ*) if *i* ≠ *j, a*_*ii*_ = −1, and *r*_*i*_ ∼ Uniform [0, 1], for different values of connectivity *C* and characteristic inter-species interaction strength *σ* > 0 (Supplementary Note S3).

Figure 2a shows the prediction error in synthetic training and test datasets, each of which has *N* samples generated by the GLV model of *N* species, with *σ* = 0.5 and different values of *C*. The prediction error in the training set, *E*(𝔇_1_), keeps decreasing with the increasing number of training epochs, especially for high *C* values (as shown in dashed and dotted cyan lines in Fig. 2a). Interestingly, the prediction error in the test dataset, *E*(𝔇_2_), reaches a plateau after enough number of training epochs regardless of the *C* values (see solid, dashed and dotted yellows lines in Fig. 2a), which is a clear evidence of an adequate training of cNODE with low overfitting. Note that the plateau of *E*(𝒟) increases with *C*. We confirm this result in datasets with different sizes of the training dataset (Fig. 2b). Moreover, we found that the plateau increases with increasing characteristic interaction strength *σ* (Fig. 2c). Fortunately, the increase of *E*(𝔇_2_) (due to increasing *C* or *σ*) can be compensated by increasing the sample size of the training set 𝔇_1_. Indeed, as shown in Fig. 2b,c, *E*(𝔇_2_) decreases with increasing |𝔇_1_| /N.

**Figure 2:**
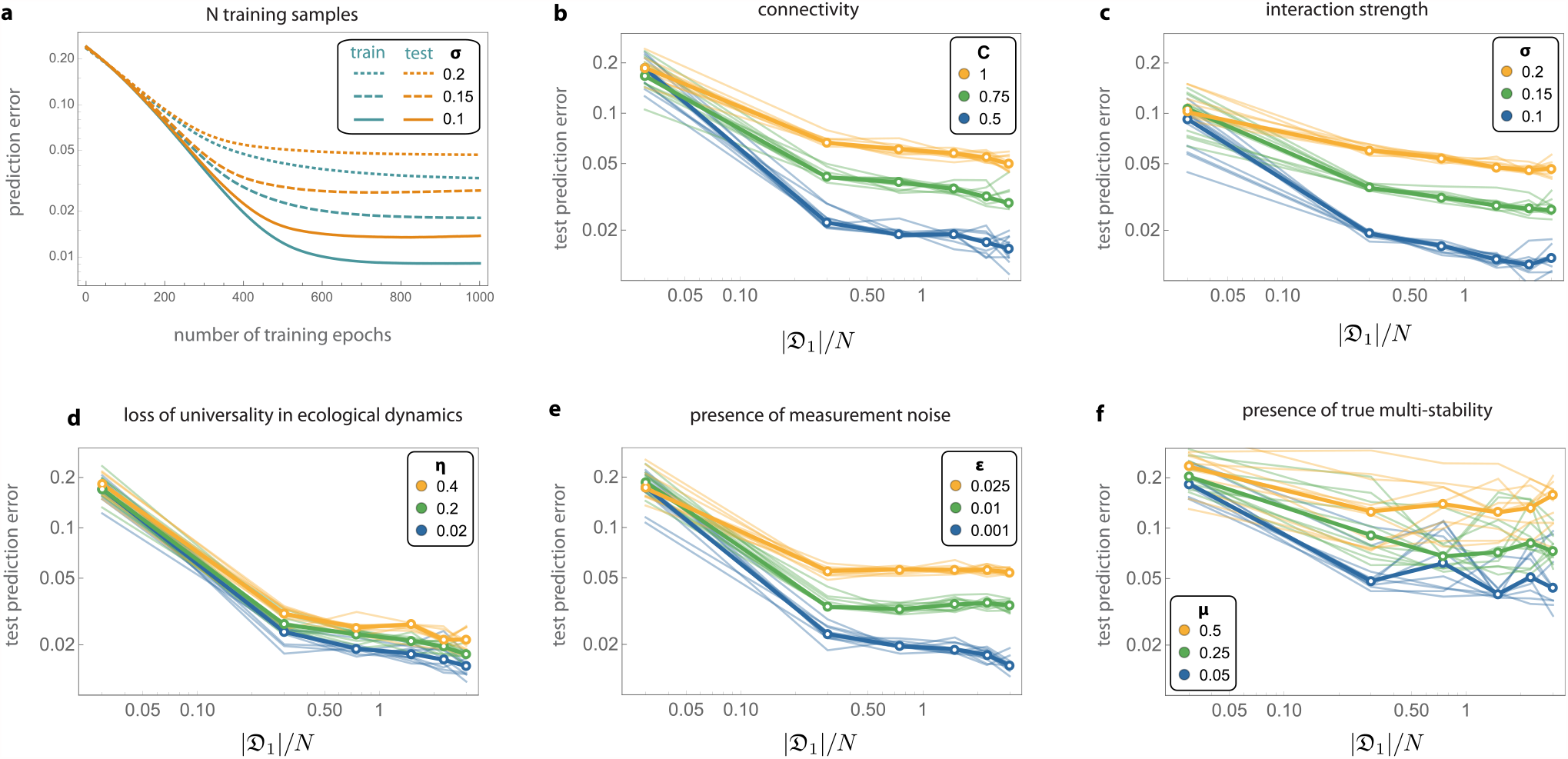
*In silico* validation of cNODE using synthetic datasets. Results are for synthetic communities of *N* =100 species generated by the with Generalized Lotka-Volterra model (panels **a**-**e**) or a population dynamics model with nonlinear functional responses (panel **f**). **a**. Training cNODE with *N* samples obtained from GLV models with connectivity *C* = 0.1 (solid), *C* = 0.15 (dashed), *C* = 0.2 (dotted). **b**. Performance of cNODE for GLV datasets with *C* = 0.5 and different interaction strengths *σ*. **c**. Performance of cNODE for GLV datasets with *σ* = 0.5 and different connectivity *C*. **d**. Performance of cNODE for GLV datasets with non-universal dynamics, quantified by the value of *η*. For all datasets, *σ* =0.1 and *C* = 0.5. **e**. Performance of cNODE for GLV datasets with measurement errors quantified by *ε*. For all datasets, *σ* = 0.1 and *C* =0.5. **f**. Performance of cNODE for synthetic datasets with multiple interior equilibria, quantified by the probability *μ* ∈ [0, 1] of finding multiple equilibria. For all datasets, *C* = 0.5, *σ* = 0.1. In panels **b**-**f**, thin lines represent the prediction errors for ten validations of training cNODE with a different dataset. Mean errors are shown in thick lines.

To systematically evaluate the robustness of cNODE against violation of its three key assumptions, we performed three types of validations. In the first validation, we generated datasets that violate the assumption of universal dynamics. For this, given a “base” GLV model with parameters (*A, r*), we consider two forms of universality loss (Supplementary Note S3). *First*, samples are generated using a GLV with the same ecological network but with those non-zero interaction strengths *a*_*ij*_ replaced by *a*_*ij*_ + Normal(0, *η*), where *η* > 0 characterizes the changes in the typical interaction strength. *Second*, samples are generated using a GLV with slightly different ecological networks obtained by randomly rewiring a proportion *ρ* ∈ [0, 1] of their edges. We find that cNODE is robust to both forms of universality loss as its asymptotic prediction error changes continuously, maintaining a reasonably low prediction error up to *η* = 0.4 and *ρ* = 0.1 (Fig. 2d and Supplementary Fig. S2).

In the second validation, we evaluated the robustness of cNODE against measurement noises in the relative abundance of species. For this, for each sample, we first change the relative abundance of the *i*-th species from *p*_*i*_ to max{0, *p*_*i*_ + Normal(0, *ε*)}, where *ε* ≥ 0 characterizes the measurement noise intensity. Then, we normalize the vector *p* to ensure it is still compositional, i.e., *p* ∈ *Δ*^*N*^. Due to the measurement noise, some species that were absent may be measured as present, and vice-versa. In this case, we find that cNODE performs adequately up to *ε* = 0.025 (Fig. 2f)

In the third validation, we generated datasets with true multi-stability by simulating a population dynamics model with nonlinear functional responses (Supplementary Notes S3). For each species collection, these functional responses generate two interior equilibria in different “regimes”: one regime with low biomass, and the other regime with high biomass. We then train cNODE with datasets obtained by choosing a fraction (1 − *μ*) of samples from the first regime, and the rest from the second regime. We find that cNODE is robust enough to provide reasonable predictions up to *μ* = 0.2 (Fig. 2d).

### Evaluation of cNODE using real data

We evaluated cNODE using six microbiome datasets of different habitats (Supplementary Note S4). The first dataset consists of *S* = 275 samples^43^ of the ocean microbiome at phylum taxonomic level, resulting in *N* = 73 different taxa. The second dataset consists of *S* = 26 *in vivo* samples of *Drosophila melanogaster* gut microbiota of *N* = 5 species^23^, as described in Fig. 1. The third dataset has *S* = 93 samples of *in vitro* communities of *N* = 8 soil bacterial species^21^. The fourth dataset contains *S* = 113 samples of the Central Park soil microbiome^22^ at the phylum level (*N* = 36 phyla). The fifth dataset contains *S* = 150 samples of the human oral microbiome from the Human Microbiome Project^24^ (HMP) at the genus level (*N* = 73 genera). The final dataset has *S* = 106 samples of the human gut microbiome from HMP at the genus level (N D 58 genera).

To evaluate cNODE, we performed the leave-one-out cross-validation on each dataset. The median test prediction errors were 0.06, 0.066, 0.079, 0.107, 0.211 and 0.242 for the six datasets, respectively (Fig. 3a). To understand the meaning of these errors, for each dataset we inspected five pairs 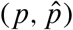 corresponding to the observed and out-of-sample predicted composition of five samples. We chose the five samples based on their test prediction error. Specifically, we selected those samples with the minimal error, close to the first quartile, closer to the median, closer to the third quartile, and with the maximal error (columns in Fig. 3b-g, from left to right). We found that samples with errors below the third quartile provide acceptable predictions (left three columns in Fig. 3b-g), while samples with errors close to the third quartile or with the maximal error do demonstrate salient differences between the observed and predicted compositions (right two columns in Fig. 3b-g). Note that in the sample with largest error of the human gut dataset (Fig. 3g, rightmost column), the observed composition is dominated by *Provotella* (pink) while the predicted sample is dominated by *Bacteroides* (blue). This drastic difference is likely due to different dietary patterns^44^.

**Figure 3:**
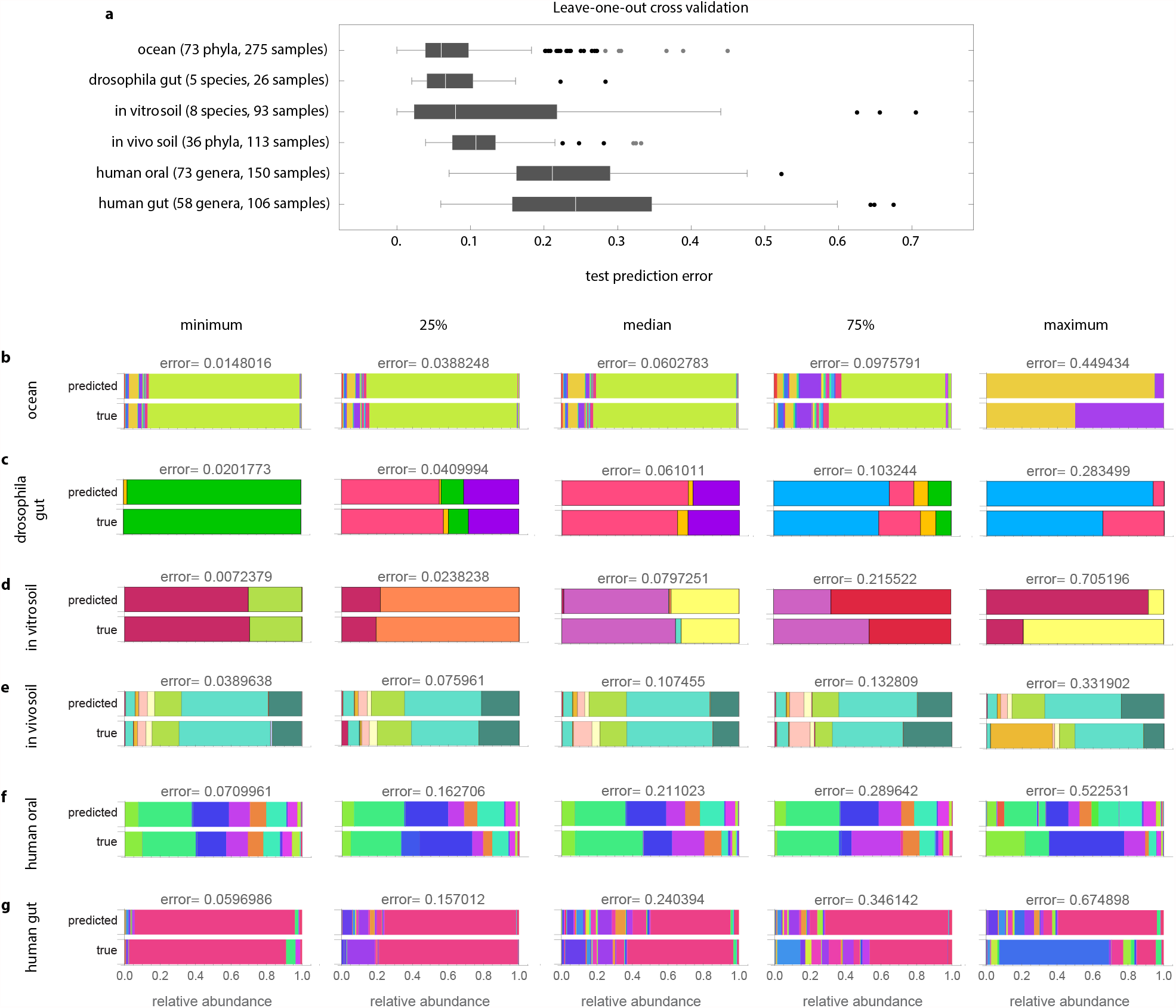
Predicting the composition of real microbiomes. **a**. Boxplots with the prediction error obtained from a leave-one-out crossvalidation of each dataset. **b**-**g**: For each dataset, we show true and predicted samples corresponding to the minimal prediction error, closer to the first quartile, median, closer to the third quartile, maximum prediction error (including outliers). Note all shown in panels **b**-**g** predictions are out-of-sample predictions.

## Discussion

cNODE is a deep learning framework to predict microbial compositions from species assemblages only. We validated its performance using *in silico, in vitro*, and *in vivo* microbial communities, finding that cNODE learns to perform accurate out-of-sample predictions using a few training samples. Classic methods for predicting species abundances in microbial communities require inference based on population dynamics models^21,41,45,46^. However, these methods typically require high-quality time-series data of species absolute abundances, which can be difficult and expensive to obtain for *in vivo* microbial communities. cNODE circumvents needing absolute abundances or time-series data. However, compared to the classic methods, the cost to pay is that the trained function *f*_*θ*_ cannot be mechanistically interpreted because of the lack of identifiability inherent to compositional data^47,48^. We also note a recent statistical method to predict the steady-state abundance in ecological communities^49^. This method also requires absolute abundance measurements. cNODE can outperform this statistical method despite using only relative abundances (Supplementary Note S6). See also Supplementary Note S5 for a discussion of how our framework compares to other related works.

Deep learning techniques are actively applied in microbiome research^50–58^, such as for classifying samples that shifted to a diseased state^59^, predicting infection complications in immunocompromised patients^60^, or predicting the temporal or spatial evolution of certain species collection^61,62^. However, to the best of our knowledge, the potential of deep learning for predicting the effect of changing species assemblages was not explored nor validated before. Our proposed framework, based on the notion of neural ODE^34^, is a baseline that could be improved by incorporating additional information. For example, incorporating available environmental information such as pH, temperature, age, BMI, and host’s diet could enhance the prediction accuracy. This additional information would help to predict the species present in different environments. Adding “hidden variables” such as the unmeasured total biomass or unmeasured resources to our ODE will enhance the expressivity of the cNODE^63,64^, but this may result in more challenging training. Finally, if available, knowledge of the genetic similarity between species can be leveraged into the loss function by using the phylogenetic Wasserstein distance^65^ that provides a well-defined gradient^66^.

We anticipate that a useful application of our framework is to predict if by adding some species collection to a local community we can bring the abundance of target species below the practical extinction threshold. Thus, given a local community containing the target (and potentially pathogenic) species, we could use a greedy optimization algorithm to identify a minimal collection of species to add such that our architecture predicts that they will decolonize the target species.

Our framework does have limitations. For example, cNODE cannot accurately predict the abundance of taxa that have never been observed in the training dataset. Also, a limitation of our current architecture is that it assumes that true multistability does not exist —namely, a community with a given species assemblage permits only one stable steady-state, where each species in the collection has a positive abundance. For complex microbial communities such as the human gut microbiota, the highly personalized species collections make it very difficult to decide if true multistability exists or not. Our framework could be extended to handle multistability by predicting a probability density function for the abundance of each species. In such a case, true multistability would correspond to predicting a multimodal density function.

In conclusion, the many species and the complex, uncertain dynamics that microbial communities exhibit have been fundamental obstacles in our ability to learn how they respond to alterations, such as removing or adding species. Moving this field forward may require losing some ability to interpret the mechanism behind their response. In this sense, deep learning methods could enable us to rationally manage and predict complex microbial communities’ dynamics.

## Acknowledgments

M.T.A. gratefully acknowledges the financial support from CONACyT project A1-S-13909, México. Y.-Y.L. acknowledges the funding support from National Institutes of Health (R01AI141529, R01HD093761, RF1AG067744, UH3OD023268, U19AI095219, and U01HL089856).

## Author contributions

M.T.A. and Y.-Y. L. conceived and designed the project. S.M.M. did the numerical analysis. S.M.M. and X.-W.W. performed the real data analysis. All authors analyzed the results. M.T.A. and Y.-Y.L. wrote the manuscript. S.M.M. and X.-W.W. edited the manuscript.

## Data accessibility

The data and code used in this work are available from the corresponding authors upon reasonable request.

