## Supplementary Notes for "Predicting microbiome compositions from species assemblages through deep learning"

**S5 Related work.** **10**

**S6 Comparing cNODE with Maynard et al.'s method.** **12**

### S1. Implementation of the compositional Neural Ordinary Differential Equation

#### S1.1 Flux implementation of cNODE.

We implemented cNODE using `Flux`[67], a library for machine learning in the Julia programming language with support for Neural Ordinary Differential Equations[34]. A complete implementation of cNODE is given in the file `cNODE.jl`.

Our implementation is based on a structure called `FitnessLayer` that contains cNODE's parameter $\Theta \in \mathbb{R}^{N\times N}$. This parameter is initialized using the Xavier's method[68]. When evaluated in a composition $p$, the `FitnessLayer` computes first $f_\theta(p) = \Theta p$. More complex functions $f_\theta$ can be easily incorporated in the code. Finally, the structure uses $f_\theta$ to calculate the right-hand side of the ODE in Eq. (2).

To predict the composition $\hat{p} \in \Delta^N$ associated with a species collection $z \in \{0,1\}^N$, cNODE numerically solves such an the ODE in Eq.(2). In `Flux`, this is automated by the function `neural_ode`, which constructs the cNODE by building a NODE with the dynamics specified by the `FitnessLayer`.

The dynamics is numerically integrated over the interval $\tau \in [0, \tau_c]$ using the `Tsit5` method[69], which is the default integration method for nonstiff ODEs in Julia. We choose $\tau_c = 1$ without loss of generality, as $\tau$ in Eq. (2) can be rescaled by multiplying $f_\theta$ by a constant. After integration, the final value at time $\tau = 1.0$ is returned as the prediction of cNODE.

The loss function calculates the average Bray-Curtis dissimilarity between there true composition $p \in \Delta^N$ and the prediction $\hat{p} \in \Delta^N$ generated by cNODE.

#### S1.2 Training cNODE.

To train cNODE, we adjust the parameters $\theta$ to minimize the loss over the training set. We experimented with two training algorithms:

1. **The ADAM algorithm**[70]. ADAM is a widely used gradient-based stochastic optimization algorithm that often compares favorably to other gradient optimization algorithms[71]. We refer to [70, 71] for additional details.

2 **ADAM plus a first-order gradient-based meta-learning algorithm based on Reptile**[37]. In the meta-learning framework, lets consider a set $\mathcal{T}$ of different *tasks* that the network needs to perform, such as learning to predict in different datasets. Reptile works by sampling some task $\tau \in \mathcal{T}$, training on it to obtain the weights $\tilde{\theta}$, and then updating the initial weights $\theta$ towards $\tilde{\theta}$.

   To train cNODE, we used ADAM plus Reptile training[37]. More precisely, we defined each task as a random partition of the training set in mini-batches. In this way, training enhances cNODE's generalizing ability from the predictions regardless of any specific partition or any sequence order of mini-batches. The algorithm we used is described below:

---

**Algorithm 1:** ADAM + Reptile for cNODE

---

Initialize $\theta$, the vector of initial parameters
**for** *epoch = 1,2...* **do**
  Random partition of training set $\rho$
  Compute $\tilde{\theta} = \text{ADAM}^k_\rho(\theta)$, denoting $k$ steps
  Update $\theta \leftarrow \theta + \text{ADAM}(\tilde{\theta} - \theta)$
**end**

---

Our implementation in the Julia language of the above algorithm is based on the model zoo of the library Flux[72].

### S2. Comparison of two deep learning frameworks in predicting microbiome compositions

#### S2.1 A ResNet architecture for predicting microbiome compositions.

We tested the performance the classical ResNet architecture[34] for predicting microbiome compositions. More precisely, we used the input layer

$$h_0 = W_0 z + b_0,$$

where $W_0 \in \mathbb{R}^{N \times N}$ and $b_0 \in \mathbb{R}^N$ are parameters to adjust. We used $L = 3$ hidden layers of the form:

$$h_\ell = h_{\ell-1} + \text{ReLU}(W_{\ell-1} h_{\ell-1} + b_{\ell-1}), \quad \ell = 1, \cdots, L,$$

where $W_\ell \in \mathbb{R}^{N \times N}$ and $b_\ell \in \mathbb{R}^N$ are parameters to adjust. Finally, the output layer takes the form

$$\hat{p} = \left( \frac{h_{L,1} z_1}{\sum_j h_{L,j} z_j}, \cdots, \frac{h_{L,N} z_N}{\sum_j h_{L,N} z_N} \right).$$

A `Flux` implementation of this ResNet can be found in the file `ResNet.jl`.
We trained this ResNet architecture using the two methods described in S1.2.

#### S2.2 Performance comparison.

We compared the performance of cNODE and ResNet architecture for predicting the composition of four out of six real microbiome datasets described in Supplementary Note S4. These datasets contain a small set of species (between 5 and 58) and samples (between 26 and 113), allowing us to make computationally expensive leave-one-out cross validation analysis. Here, we also compared the effect of training both architectures with ADAM, and with ADAM plus the Reptile like metalearning algorithms.

To perform the comparison we used a leave-one-out cross validation on each of the four datasets (Supplementary Fig. S1). From these results, some remarks are in order:

1. For both *in-vivo* datasets, cNODE trained with Reptile outperforms all other algorithms.

2. The ResNet architecture trained using only ADAM can provide reasonably accurate prediction for simple small species collections (i.e., for the $N = 5$ species in Drosophila gut and the $N = 8$ species in the soil *in-vitro* community).

3. For cNODE, training using the Reptile metalearning algorithm decreases the prediction error in the test dataset. Interestingly, using Reptile does not alway decreases the prediction error in the train dataset. Therefore, the Reptile metalearning algorithm is performing as desired to enhance the generalizing ability of cNODE.

4. For ResNet, training with the Reptile metalearning algorithm can increase the prediction errors in both the training and test datasets when compared to training only with ADAM.

5. The ResNet architecture exhibits a higher variability in the training set when compared to cNODE. This suggests that the performance of ResNet is significantly influenced by the initialization parameters. In particular, training a ResNet with Reptile can significantly increase the variability of prediction errors (see, e.g., the soil *in vitro* dataset).

Overall, the above remarks indicate that cNODE training with Reptile outperforms the other architectures when predicting complex microbial communities like human gut microbiota or *in vivo* soil communities.

### S3. Population dynamics for the *in silico* validations.

We generated *in-silico* species pools using the Generalized Lotka-Volterra (GLV) equations, a classical population dynamics model successfully applied to diverse microbial communities, from soils[39] and cheese[45], to the human body[11,41]. The GLV model takes the form

$$\frac{\mathrm{d}x(t)}{\mathrm{d}t} = x(t) \odot [Ax(t) + r], \quad x(0) = x_0, \tag{S1}$$

where $x(t) = (x_1(t), \cdots, x_N(t))^\top \in \mathbb{R}_{\geq 0}$ and $x_i(t)$ denotes the absolute abundance of species $i$ at time $t$. Above, $x \odot v$ is the entry-wise multiplication of vectors $x, v \in \mathbb{R}^N$. The GLV model has two parameters: the interaction matrix of the species pool $A = (a_{ij}) \in \mathbb{R}^{N\times N}$, and the intrinsic growth-rate of the species $r = (r_i) \in \mathbb{R}^N$. In particular, the $j$-th species has a positive impact on the $i$-th species if $a_{ij} > 0$, a negative impact if $a_{ij} < 0$, and no impact if $a_{ij} = 0$. Recall also that the interaction matrix determines the underlying ecological network $\mathcal{G}(A)$ of the species pool. This network has one vertex associated to each species and an edge $(j \to i) \in \mathcal{G}(A)$ if $a_{ij} \neq 0$.

To generate the relative abundance vector $p \in \Delta^N$ corresponding to a local community with species collection $z \in \{0, 1\}^N$, we follow four steps:

1. Set the parameters $(A, r)$.
2. Set the initial abundance of species $x_0 \in \mathbb{R}^N_{\geq 0}$ as

   $$x_{0,\,i} = \begin{cases} 0 & \text{if } z_i = 0, \\ \text{Uniform}[0, 1] & \text{otherwise,} \end{cases}$$

   for $i = 1, \cdots, N$.
3. Numerically integrate Eq. (S1) with initial condition $x_0$ until the system reaches a steady-state abundance $x^* = (x^*_1, \cdots, x^*_N) \in \mathbb{R}^N$. For the results presented in our paper, we choose a final integration time $t_f = 1000$.
4. Compute the relative abundance vector $p = (p_1, \cdots, p_N)^T \in \Delta^N$ as $p_i = x^*_i / \sum_j x^*_j$.

Using the above procedure, we generated a dataset $\mathfrak{D}$ by randomly sampling species collections $z \in \{0, 1\}^N$ and calculating the corresponding $\hat{p} \in \Delta^N$. Below we detail the construction of the three types of datasets used in the Main Text.

#### S3.1 Generating datasets with universal dynamics.

To generate a dataset with universal dynamics, we considered that all species collections have the same parameters $(A, r)$. These parameters were generated as follows. The interaction matrix $A$ of the community is obtained as the adjacency matrix of a directed weighted Erdös-Rényi random network with connectivity $C \in [0, 1]$. The edge-weights were chosen from a Normal distribution with zero mean and variance $\sigma^2$, where $\sigma > 0$ represents the "characteristic" inter-species interaction strength. The intrinsic growth $r_i$ is chosen uniformly at random from the interval [0, 1].

#### S3.2 Generating datasets with non-universal dynamics.

To generate a dataset with non-universal dynamics, we considered two possible sources for non-universality. First, the mechanisms of interaction between species may differ across local communities. In Eq. (S1), this translates as using different parameters $a_{ij}$ for each non-zero interaction in different local communities. Thus, in this case we replaced each $a_{ij} \neq 0$ by $a_{ij} + \eta\text{Normal}(0, 1)$, where $\eta > 0$ quantifies the *changes in the typical interaction strength*, and hence the "loss" of universality in this case.

Second, we considered that each local community may have a different ecological network. To model this case, we considered that the ecological network of each local community is obtained by *randomly rewiring* a proportion $\rho \in [0, 1]$ of the edges of a baseline ecological network $\mathcal{G}(A)$, thus shuffling a proportion of entries of the associated $A$ matrix. Since $\rho = 0$ corresponds to universal dynamics, the magnitude of $\rho$ quantifies the "loss" of universality in this case.

#### S3.3 Adding measurement noise to a dataset.

For a pair $(z, p)$ in a dataset $\mathfrak{D}$, we added noise by replacing $p_i$ by first adding a small noise $w_i = \max\{0, p_i + \varepsilon\text{Normal}(0, 1)\}$, and then normalizing to obtain the noisy measurement $p_i \leftarrow w_i / \sum_j w_j$. Here, the parameter $\varepsilon > 0$ controls the *measurement noise intensity*.

#### S3.4 Generating datasets with multi-stability.

To generate a dataset with true multi-stability, we calculated the steady-states from a population dynamics model with the following non-linear functional response:

$$\frac{\mathrm{d}x_i(t)}{\mathrm{d}t} = x_i(t)\left[r_i + \sum_{j=1}^{N} a_{ij}\,\frac{x_j(t)}{1 + h\,x_j(t)^2}\right], \quad i = 1, \cdots, N, \tag{S2}$$

where $h$ denotes the handling time.

To generate steady-states with multi-stability, we first select a GLV model with a linear functional response (Eq. S1) and universal dynamics (Supplementary Notes S3), and compute the steady-state $\xi^*$. Note that the steady-state abundances satisfies the equation

$$r_i = -\sum_{j=1}^{N} a_{ij}\,\xi_j^*. \tag{S3}$$

The steady-states of Eq.(S2) satisfies the equation

$$\sum_{j=1}^{N} a_{ij}\,\frac{x_j}{1 + h\,x_j^2} + r_i = 0, \tag{S4}$$

so that we substitute Eq.S3 in Eq.S4 and solve for $x_j$ the following quadratic equation:

$$h\xi_j^* x_j^2 - x_j + \xi_j^* = 0, \tag{S5}$$

for all $j$, and then we compute the relative abundance vector $p = (p_1, \cdots, p_N)^T \in \Delta^N$ as $p_i = x_i^* / \sum_j x_j^*$. To ensure that there are two real solutions for $x_j$, we chose $h = \frac{1}{4\xi_k^{*2}} > 0$ for some $k$.

The two steady-state abundances corresponds to a high and low total biomass regimes, respectively. To build the datasets, we chose a fraction (1-$\mu$) from the first regime, and the rest from the second.

### S4. Description of the experimental datasets.

#### S4.1 In-vivo drosophila core gut microbiome.

The drosophila dataset[23] contains the absolute abundance of the five species in each possible local community with different species collection. See Supplementary Table S1 for species IDs. There is five replicates for each of those species collections. We averaged those five replicates, discarded samples with a single species, and obtained the relative abundance of each of the remaining samples. This yielded 31 samples with different species collection.

| ID | Genus | Species |
|---|---|---|
| 1 | Lactobacillus | plantarum |
| 2 | Lactobacillus | brevis |
| 3 | Acetobacter | pasteurianus |
| 4 | Acetobacter | tropicalis |
| 5 | Acetobacter | orientalis |

Supplementary Table S1: Species IDs Drosophila gut microbiota.

#### S4.2 In-vitro soil community.

This laboratory community of $N = 8$ heterotrophic soil-dwelling bacterial species[21] described in Table S2. The available dataset contains 98 samples, including all solos, all duos, some trios, one septet and one octet.

| ID | Genus | Species |
|---|---|---|
| 1 | Enterobacter | aerogenes |
| 2 | Pseudomonas | aurantiaca |
| 3 | Pseudomonas | chlororaphis |
| 4 | Pseudomonas | citronellolis |
| 5 | Pseudomonas | fluorescens |
| 6 | Pseudomonas | putida |
| 7 | Pseudomonas | veronii |
| 8 | Serratia | marcescens |

Supplementary Table S2: Species IDs the in-vitro soil community.

#### S4.3 In-vivo soil microbiome.

The soil dataset consists of soil microbiome across Central Park in New York City consist of 1160 samples. This data set is 16S rRNA gene-based with variable region V4. The data is available at https://qiita.ucsd.edu/ under study ID 2140 and the detailed description of this data set can be found in Ref. [22]. We used the function `summarize_taxa.py` QIIME 1 to summarize taxa to different taxonomic levels with defaults options. Supplementary Table S3 provides the IDs associated to each phylumm.

#### S4.4 Human gut microbiome.

A 16S rRNA gene-based data set from variable regions V3 to V5. The data are available at http://www.hmpdacc.org/HMQCP/. We selected the samples from the stool body site. For multiple samples from the same subject, we only keep one single sample of that subject. To guarantee the

| ID | Kindgom | Phylum |
|---|---|---|
| 1 | Archaea | Crenarchaeota |
| 2 | Archaea | Euryarchaeota |
| 3 | Archaea | Parvarchaeota |
| 4 | Bacteria | others |
| 5 | Bacteria | Acidobacteria |
| 6 | Bacteria | Actinobacteria |
| 7 | Bacteria | Aquificae |
| 8 | Bacteria | Armatimonadetes |
| 9 | Bacteria | BHI80-139 |
| 10 | Bacteria | Bacteroidetes |
| 11 | Bacteria | Chlorobi |
| 12 | Bacteria | Chloroflexi |
| 13 | Bacteria | Cyanobacteria |
| 14 | Bacteria | Elusimicrobia |
| 15 | Bacteria | FBP |
| 16 | Bacteria | Firmicutes |
| 17 | Bacteria | GN02 |
| 18 | Bacteria | Gemmatimonadetes |
| 19 | Bacteria | Lentisphaerae |
| 20 | Bacteria | NC10 |
| 21 | Bacteria | Nitrospirae |
| 22 | Bacteria | OD1 |
| 23 | Bacteria | OP3 |
| 24 | Bacteria | OP9 |
| 25 | Bacteria | Planctomycetes |
| 26 | Bacteria | Proteobacteria |
| 27 | Bacteria | SBR1093 |
| 28 | Bacteria | Spirochaetes |
| 29 | Bacteria | TM6 |
| 30 | Bacteria | TM7 |
| 31 | Bacteria | TPD-58 |
| 32 | Bacteria | Tenericutes |
| 33 | Bacteria | Verrucomicrobia |
| 34 | Bacteria | WPS-2 |
| 35 | Bacteria | WS1 |
| 36 | Bacteria | WS6 |

Supplementary Table S3: Phylum IDs for soil dataset.

model can be trained sufficiently, we summarized the taxa into the genus level and removed the genus with fewer than 50 reads. See also Supplementary Table S4 for genus ID.

| ID | Phylum | Class | Order | Family | Genus |
|---|---|---|---|---|---|
| 1 | Firmicutes | Clostridia | Clostridiales | Veillonellaceae | *Veillonella* |
| 2 | Firmicutes | Clostridia | Clostridiales | Ruminococcaceae | *Clostridium* |
| 3 | Firmicutes | Clostridia | Clostridiales | Ruminococcaceae | *Bacteroides* |
| 4 | Tenericutes | Erysipelotrichi | Erysipelotrichales | Erysipelotrichaceae | *Coprobacillus* |
| 5 | Firmicutes | Clostridia | Clostridiales | ClostridialesFamilyXIII.IncertaeSedis | |
| 6 | Bacteroidetes | Bacteroidia | Bacteroidales | Porphyromonadaceae | *Odoribacter* |
| 7 | Firmicutes | Clostridia | Clostridiales | Lachnospiraceae | *Lachnobacterium* |
| 8 | Verrucomicrobia | Verrucomicrobiae | Verrucomicrobiales | Verrucomicrobiaceae | *Akkermansia* |
| 9 | Bacteroidetes | | | | |
| 10 | Proteobacteria | Gammaproteobacteria | Pasteurellales | Pasteurellaceae | *Haemophilus* |
| 11 | Firmicutes | Clostridia | Clostridiales | Veillonellaceae | *Megasphaera* |
| 12 | Firmicutes | Bacilli | Lactobacillales | Streptococcaceae | *Streptococcus* |
| 13 | Firmicutes | Clostridia | Clostridiales | Ruminococcaceae | *Anaerotruncus* |
| 14 | Bacteroidetes | Bacteroidia | Bacteroidales | Porphyromonadaceae | *Parabacteroides* |
| 15 | Firmicutes | Clostridia | Clostridiales | Dehalobacteriaceae | *Dehalobacterium* |
| 16 | Tenericutes | | | | |
| 17 | Firmicutes | Clostridia | Clostridiales | Lachnospiraceae | *Ruminococcus* |
| 18 | Firmicutes | Clostridia | Clostridiales | Lachnospiraceae | *Dorea* |
| 19 | Tenericutes | Erysipelotrichi | Erysipelotrichales | Erysipelotrichaceae | *Catenibacterium* |
| 20 | Proteobacteria | Deltaproteobacteria | Desulfovibrionales | Desulfovibrionaceae | *Desulfovibrio* |
| 21 | Firmicutes | Clostridia | Clostridiales | Ruminococcaceae | *Oscillospira* |
| 22 | Firmicutes | Clostridia | Clostridiales | Lachnospiraceae | *Clostridium* |
| 23 | Proteobacteria | Gammaproteobacteria | Enterobacteriales | Enterobacteriaceae | *Escherichia* |
| 24 | Firmicutes | Bacilli | Turicibacterales | Turicibacteraceae | |
| 25 | Firmicutes | Clostridia | Clostridiales | Ruminococcaceae | |
| 26 | Actinobacteria | Actinobacteria | Coriobacteriales | Coriobacteriaceae | *Collinsella* |
| 27 | Firmicutes | Clostridia | Clostridiales | Veillonellaceae | *Acidaminococcus* |
| 28 | Firmicutes | Clostridia | Clostridiales | ClostridialesFamilyXIII.IncertaeSedis | *Eubacterium* |
| 29 | Firmicutes | Bacilli | Lactobacillales | Lactobacillaceae | *Lactobacillus* |
| 30 | Firmicutes | Clostridia | Clostridiales | Lachnospiraceae | *Roseburia* |
| 31 | Firmicutes | Clostridia | Clostridiales | Ruminococcaceae | *Eubacterium* |
| 32 | Firmicutes | Clostridia | Clostridiales | Lachnospiraceae | *Lachnospira* |
| 33 | Bacteroidetes | Bacteroidia | Bacteroidales | | |
| 34 | Proteobacteria | Deltaproteobacteria | Desulfovibrionales | Desulfovibrionaceae | *Bilophila* |
| 35 | Bacteroidetes | Bacteroidia | Bacteroidales | Prevotellaceae | *Prevotella* |
| 36 | Bacteroidetes | Bacteroidia | Bacteroidales | Rikenellaceae | *Alistipes* |
| 37 | Firmicutes | Clostridia | Clostridiales | Veillonellaceae | *Dialister* |
| 38 | Firmicutes | Clostridia | Clostridiales | Ruminococcaceae | *Ruminococcus* |
| 39 | Proteobacteria | Betaproteobacteria | Burkholderiales | | |
| 40 | Tenericutes | Erysipelotrichi | Erysipelotrichales | Erysipelotrichaceae | *Holdemania* |
| 41 | Fusobacteria | Fusobacteria | Fusobacteriales | Fusobacteriaceae | *Fusobacterium* |
| 42 | Firmicutes | Clostridia | Clostridiales | Lachnospiraceae | *Eubacterium* |
| 43 | Firmicutes | Clostridia | Clostridiales | Ruminococcaceae | *Subdoligranulum* |
| 44 | Firmicutes | Clostridia | Clostridiales | Ruminococcaceae | *Faecalibacterium* |
| 45 | Tenericutes | Erysipelotrichi | Erysipelotrichales | Erysipelotrichaceae | *Clostridium* |
| 46 | Proteobacteria | Betaproteobacteria | Burkholderiales | Alcaligenaceae | *Sutterella* |
| 47 | Others | | | | |
| 48 | Firmicutes | Clostridia | Clostridiales | Clostridiaceae | *Clostridium* |
| 49 | Firmicutes | Clostridia | Clostridiales | Lachnospiraceae | *Coprococcus* |
| 50 | Firmicutes | Clostridia | Clostridiales | Lachnospiraceae | *Blautia* |
| 51 | Bacteroidetes | Bacteroidia | Bacteroidales | Bacteroidaceae | *Bacteroides* |
| 52 | Firmicutes | Clostridia | Clostridiales | Clostridiaceae | |
| 53 | Firmicutes | Clostridia | Clostridiales | Lachnospiraceae | *Bacteroides* |
| 54 | Firmicutes | Clostridia | Clostridiales | Veillonellaceae | *Phascolarctobacterium* |
| 55 | Firmicutes | Clostridia | Clostridiales | | |
| 56 | Actinobacteria | Actinobacteria | Bifidobacteriales | Bifidobacteriaceae | *Bifidobacterium* |
| 57 | Firmicutes | Clostridia | Clostridiales | Veillonellaceae | *Megamonas* |
| 58 | Firmicutes | Clostridia | Clostridiales | Lachnospiraceae | |

Supplementary Table S4: Genus IDs for the human gut microbiota dataset.

### S5. Related work.

Here we describe related methods, emphasizing some key differences with respect to our framework.

1. **Abundance prediction based on inferred population dynamics.** A classical method to predict species abundance in microbial ecosystems is modeling their population dynamics[11,21,41,45,46]. Typically, the model is a set of parametrized ODEs —such as the Generalized Lotka-Volterra equations— describing the changes over time in the *absolute* abundances of a set of species. The model is fitted to available temporal data of absolute abundance to infer parameters, such as intrinsic growth rates, inter-species interaction strengths, etc. Then, to predict the abundance of certain species collection, the fitted ODE model is solved starting from suitable initial condition. However, applying this method for large microbial communities like the human gut is challenging if not impossible because: 1) the absence of high-quality temporal data; and 2) typically only relative abundance of species are measured. Furthermore, because the very broad population dynamics that ecosystems display even at the scale of two species[73], it is always very challenging to choose an adequate parametrized ODE model for the population dynamics of the community.

2. **Predictions based on neural networks methods.** Larsen et al.[61] employed an artificial neural network to predict the *temporal* evolution of the composition of bacterial communities with a constant species collection. More precisely, they developed a bioclimatic model of relative microbial abundance that specifically incorporates interactions between biological units. They modeled the complex interactions between microbial taxa and their environment as an artificial neural network (ANN). This method is based on two key assumptions: (1) community patterns share mathematically describable relationships with environmental conditions; and (2) the ecosystem maintains a persistent microbial community. Note that the second assumption implies that this method can not be used to predict the impact of changing the species collections. Compared to method based on inferring population dynamics, this method has the advantage of not requiring to specify any model for the community dynamics. However, in contrast to our framework, this method cannot predict the effect of changing the species collection.

   Similarly, the recent work of Zhou et al.[62] uses a neural network to predict the temporal and *spatial* evolution of the composition of microbial communities with a constant species collection. More precisely, the authors formulated the prediction of microbial communities at unsampled locations as a multi-label classification task, where each location is considered as an instance and each label represents a microbe species. Based on a set of heterogeneous features extracted from the urban environment, they aimed to predict the presence or absence of a list of microbes species at a nearby location. Note, however, that this method cannot be immediately used to predict the effect of changing the species collection.

3. **Predictions based on statistical methods.** Recently, Maynard et al.[49] proposed a statistical method to predict species abundances from species collections. More precisely, based on measuring the absolute abundance of species at some steady-states of the ecosystem, this method assumes a linear model to predict all other steady-states. For the method to be applicable, it requires that the following assumptions are satisfied: (1) each species must be present in at least $n$ distinct endpoints, not counting replicates; (2) each species must co-occur with each other species in at least one endpoint (that is, for every pair of species $i$ and $j$, there must be some endpoint where $i$ and $j$ co-occur, possibly along with other species); and (3) for each $i$ there must exist a perfect matching between the $n$ species and

the endpoints in which they co-occur with $i$. As explained in the original manuscript[49], conditions (1) and (2) above requires that "coexistence among species must be reasonably widespread for [these] conditions to hold." This method may be challenging to apply for microbial communities because it requires measuring absolute abundances. Furthermore, because microbial communities tend to have nonlinear behaviors even at the scale of two species[73,74], the implicit assumption of linearity may fail to be satisfied. Finally, we note that cNODE does not require any of the above three assumptions to be applicable, although its prediction accuracy may be influenced by them.

Similarly, Tung et al.[75] use a linear regression method to predict species compositions from information of social networks of individuals. More precisely, the authors fit a classical linear mixed model to predict the relative abundance of species in a sample based on the following predictors: social group membership, age, sex, and read depth. We note here that the predictors in this approach are completely different than the predictors used in cNODE (i.e., it only uses species collections), and thus are not comparable.

### S6. Comparing cNODE with Maynard et al.'s method.

Here we compare the performance of cNODE with the method of Maynard et al. for predicting endpoints[49]. This last method is described with details in item 3 of Supplementary Note S5.

To perform the comparison, we generated *in silico* datasets of $N = 5$ species with Generalized Lotka-Volterra dynamics. More precisely, we generated datasets with universal dynamics, maintaining the connectivity $C = 0.5$ constant and changing the typical interaction strength as $\sigma \in \{0.1, 0.2, 0.3, 0.4\}$. For each value of $\sigma$, we generated 10 datasets containing all $S = 2^5 - 1$ samples with different species collections following the simulation method described in Supplementary Note S3. We repeated this simulation method three times, obtaining three repetitions for the abundance of each species collections that can be used in Maynard's method. Additionally, because Maynard's method requires absolute abundances, we kept in the datasets both the absolute abundance and relative abundance of each steady-state that is reached. Using these datasets, we constructed training datasets by randomly choosing 70% of the samples, and the rest of the samples as test datasets.

To adjust Maynard's method we used the default parameters that were selected for $N = 4$ species. After this, we obtained the corresponding absolute abundance predictions of the test dataset. Finally, to allow a comparison with cNODE that predicts relative abundances, we transformed each predicted absolute abundance into a predicted relative abundance, and then calculated the prediction error using the Bray-Curtis dissimilarity.

For cNODE, we used the exact same training dataset as for Maynard's method, the only difference being that we trained cNODE with relative abundances. We choose a inner learning rate of 0.001, an outer learning rate of 0.005, and trained cNODE for 500 epochs using mini batches of 5 samples. We then calculated the prediction error in the test dataset using the Bray-Curtis dissimilarity. We emphasize that cNODE uses less information that Maynard's method, in the sense that the total biomass of each sample is unknown in this case.

The results of the comparison of Maynard's method and cNODE are shown in Fig. S3. We find that, for the above conditions used to construct the datasets, cNODE outperforms Maynard's method in both the training and test datasets. Crucially, note that cNODE was trained using only relative abundance measurements. We do not claim this results holds for all datasets, as there might be cases where the assumptions required by Maynard's method are exactly satisfied but none of the assumptions of cNODE are satisfied.

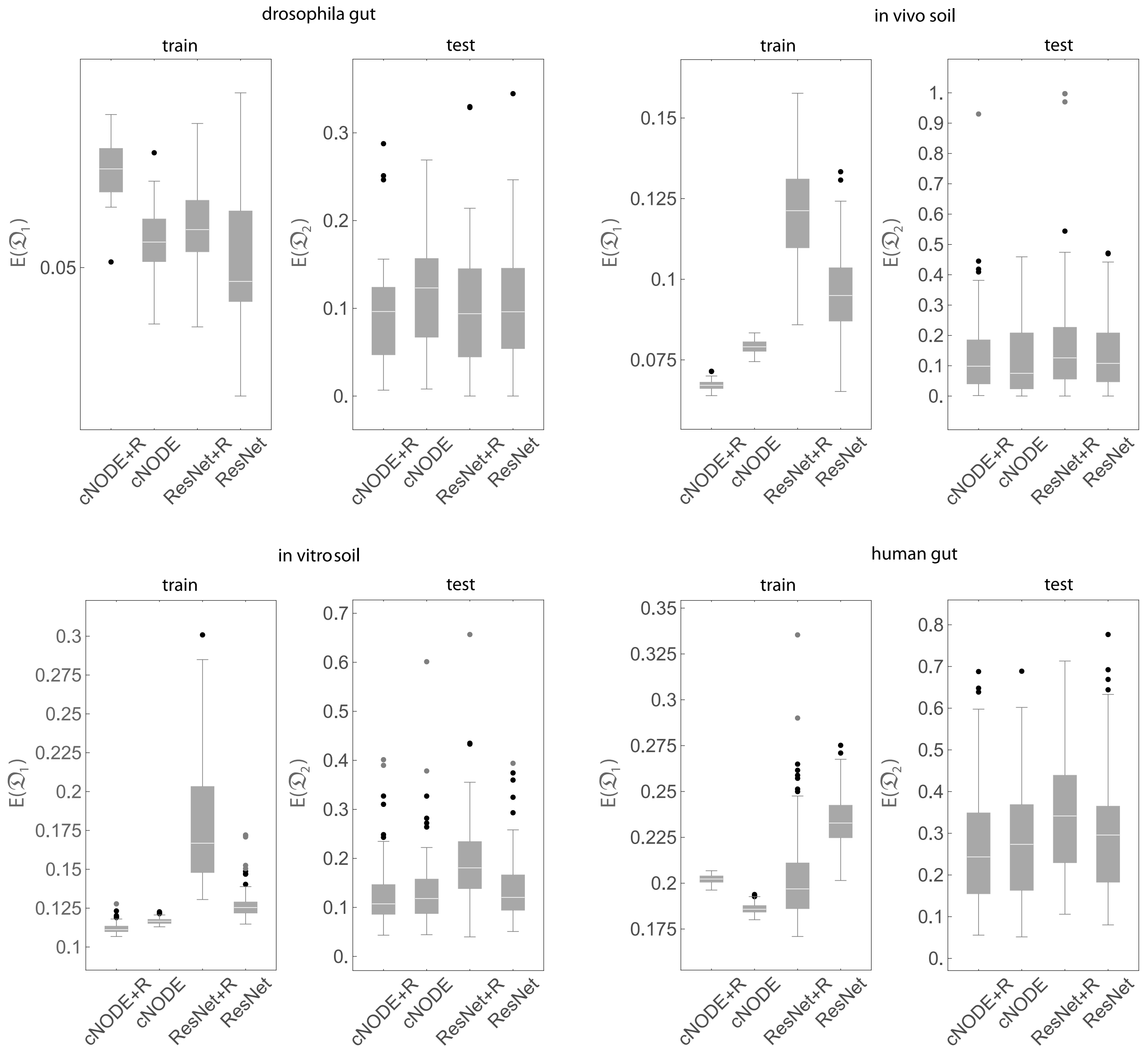

Supplementary Figure S1: **Performance of the ResNet and cNODE architectures for predicting compositions in experimental microbiomes.** Vertical axis denotes prediction error.

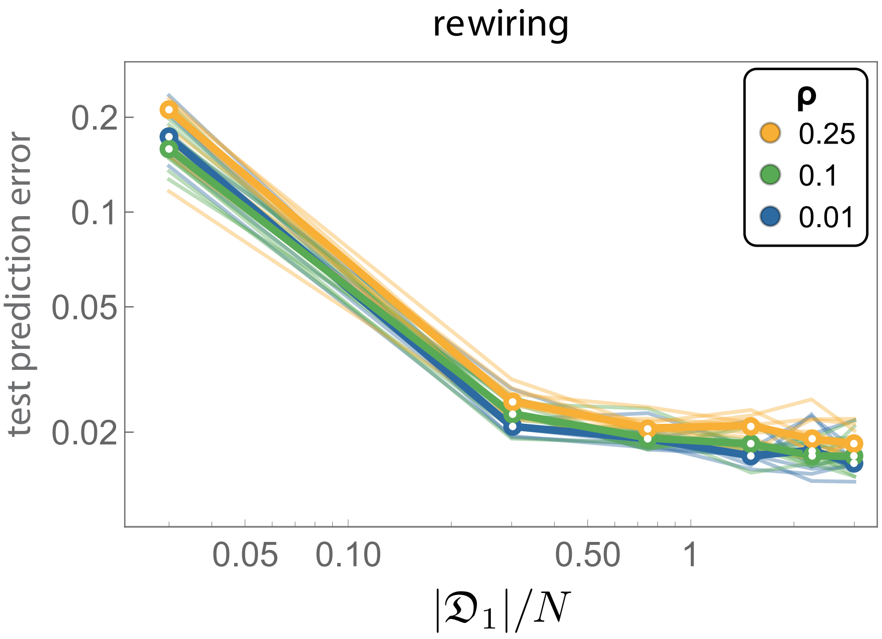

Supplementary Figure S2: **Performance of the ResNet and cNODE architectures for predicting compositions in experimental microbiomes.** Vertical axis denotes prediction error.

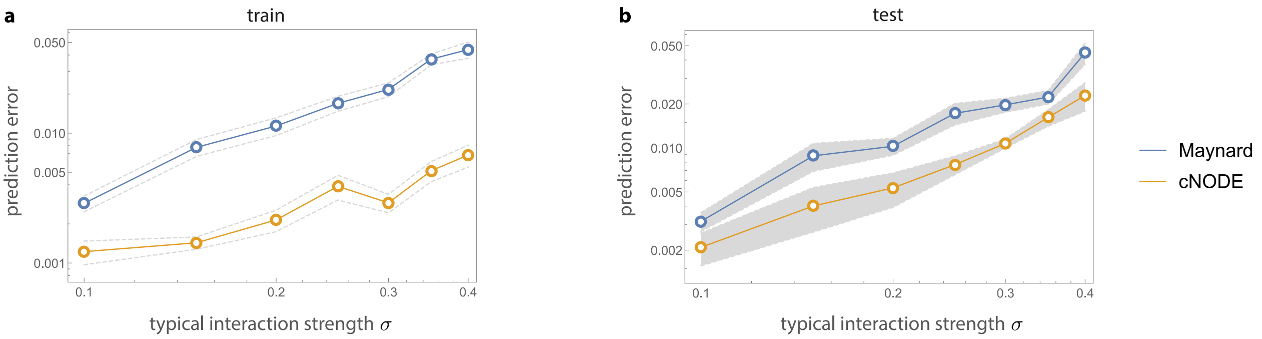

Supplementary Figure S3: **Prediction errors of Maynard et al.'s method[49] and cNODE.** For an *in silico* dataset of $N = 5$ species with universal dynamics and different typical interaction strength. Circles denote mean error for 10 repetitions, and gray shadows indicate standard deviation of the mean. **a.** Train dataset. **b.** Test dataset.
